# Enzyme-functionalized microparticles to open the vitreoretinal interface

**DOI:** 10.64898/2026.01.12.699029

**Authors:** Lucie Motyčková, Florian Peter, Ludwig Geisweid, Niklas Junker, María I. Real de Asúa Pérez-Serrano, Munisa Tabarova, Ronald Curticean, Irene Wacker, Rasmus R. Schröder, Maximilian Hammer, Dimitris Missirlis, Mariana Alarcón-Correa, Peer Fischer

**Author notes:** These authors contributed equally to this work (Lucie Motyčková, Florian Peter).

## Abstract

The transport of therapeutics and gene carriers to their site of action is often hindered by biological barriers, such as cell layers and basement membranes. Among these, the inner limiting membrane (ILM) represents a major barrier within the eye, separating the vitreous body from the retina. The ILM must be crossed, if for instance gene carriers are to reach retinal target cells following intravitreal administration. However, the ILM is a densely cross-linked basement membrane barrier, allowing only the smallest nanoparticles to pass. Here, we demonstrate that active microcolloids decorated with enzymes can locally open the ILM and thereby facilitate the diffusion of passive carriers into retinal tissue. We utilize an *ex vivo* porcine eye model to determine the membrane permeability threshold using fluorescent nanoprobes. We further show that collagenase-decorated silica microparticles can facilitate the transport of nanoparticles, while exhibiting excellent biocompatibility with no adverse morphological or functional retinal effects over a six-week *in vivo* evaluation in a porcine model. Overall, our findings introduce a biocompatible and minimally invasive strategy to facilitate the targeted nanoparticle transport across biological barriers, which we demonstrate for retinal delivery enabled by active colloids.

## 1 Introduction

In recent years, several gene therapies have been proposed^[1,2]^, but the delivery of necessary agents can be challenging as biological barriers often present a formidable hurdle. One such example is ocular gene therapy, which could target inherited retinal diseases^[1]^, a major cause of vision loss with an estimated prevalence of approximately 5.5 million people worldwide^[3]^. However, effective delivery to target affected retinal cells — such as photoreceptors and retinal pigment epithelium (RPE) cells — remains a major challenge.^[2]^ Current clinical administration of retinal gene therapy, including a recent, encouraging report in children^[4]^, still rely on viral vector delivery via complex subretinal injections. This surgical technique delivers vectors between the photoreceptors and the underlying RPE^[2]^, leading to partial retinal detachment and increasing the risk of severe complications such as permanent retinal detachment or hole formation.^[5]^ Moreover, the treatment is confined to the immediate vicinity of the injection site.^[6]^ To overcome these limitations, intravitreal injection is being explored as a less invasive alternative.^[7,8]^

The efficiency of gene delivery through intravitreal administration is significantly hindered by biological barriers, particularly the inner limiting membrane (ILM) — a basement membrane of the Müller glia at the vitreoretinal interface composed of extracellular matrix proteins, such as collagen IV, laminin, and proteoglycans.^[9,10]^ These components form a dense three-dimensional network that impedes the penetration of nanoscale gene carriers in the retina and hence their ability to reach target cells. The estimation of the ILM size cut-off in humans has been investigated with various *in vitro* and *ex vivo* animal models, but remains poorly understood due to multiple factors: i) lack of *in vivo* data owing to accessibility issues, ethical burden and high cost, ii) minimal ability to translate existing *in vitro*/*ex vivo* results to the *in vivo* context^[11]^; iii) post-mortem cessation of physiological processes and consequent rapid retina degradation during *ex vivo* studies^[12,13]^; iv) species-dependent differences in the ILM structure.^[14]^ Collectively, *ex vivo* experimentation emphasize the use of the most representative animal models - favoring species more closely related to humans over rodents -^[11]^ and advocate for minimizing the time elapsed post-mortem to preserve physiological relevance.^[15]^ A review of investigations in large animal models employing particles, FITC-dextrans, antibodies, and lipoplexes indicate that there is a size-exclusion limit of the ILM^[16–23]^. Of note, most studies evaluated the permeability of negatively charged molecules or nanoparticles, which should limit interactions with the predominantly negatively charged macromolecules in the vitreous.^[24–26]^ There is however no consensus on the exact size limit, which has been reported to lie between 2 and 20 nm^[16–20]^, with a few reports suggesting thresholds even above 40 nm.^[21–23]^ These sizes are highly relevant when considering potential transfection agents. For example, adeno-associated viral vectors, some of the smallest, widely-employed transfection agents for retinal gene therapy have capsids approximately 22-25 nm in diameter.^[27,28]^ Likewise, non-viral alternatives, such as compacted DNA-peptide nanoparticles have been engineered within a comparable size range (tens of nanometers in diameter), depending on the gene payload and particle conformation.^[29]^ Given that the physical dimensions of these available delivery vectors approach or exceed the estimated ILM size threshold, unaided passive diffusion across the membrane appears unlikely or at best challenging, highlighting the need for novel, advanced strategies that ensure the efficient transport of therapeutic genes to target retinal cells.

Here, we first use fluorescent nanoprobes in a newly developed ex-vivo porcine model to assess the permeability of the ILM (Fig. 1**A**). We then show that microparticles decorated with collagenase enable the localized enzymatic degradation of the ILM (Fig. 1**B**), thereby facilitating the diffusion of 25 nm particles into deeper retinal layers (Fig. 1**C**). Finally, we demonstrate the biocompatibility of collagenase-functionalized particles *in vivo* through intravitreal injections into the eyes of Landrace pigs as a relevant large-animal model, thereby supporting their translational potential for enhanced retinal gene delivery.

**Figure 1:**
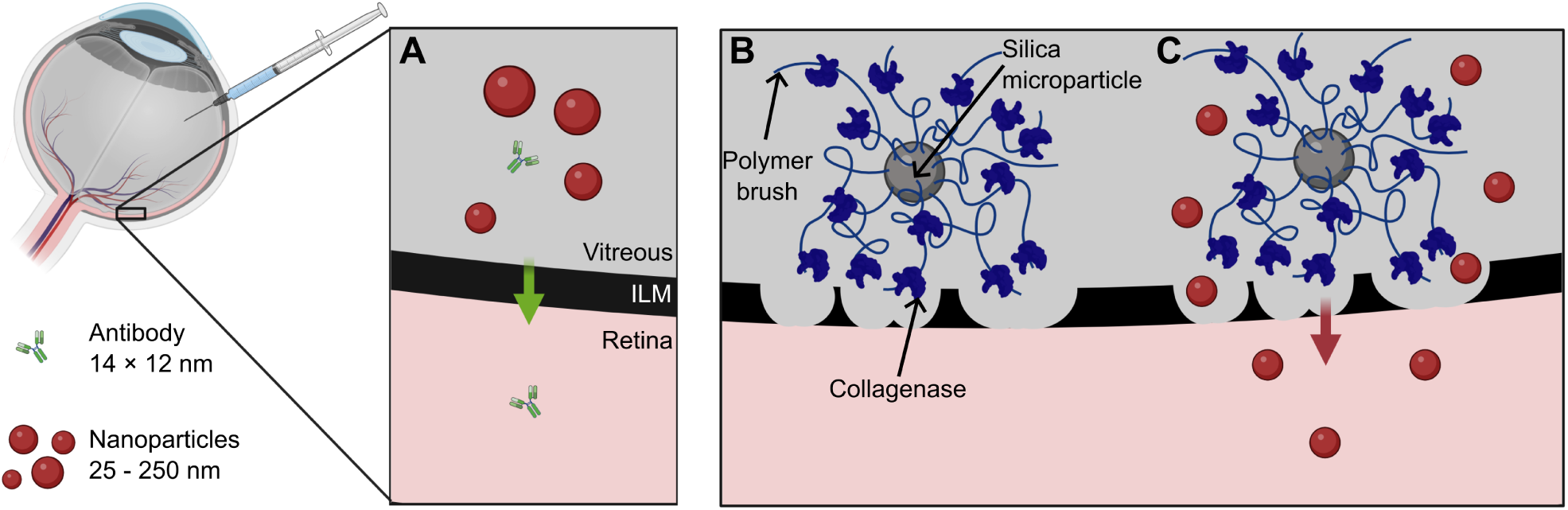
Schematic illustration of nanomaterial traversal across the inner limiting membrane (ILM): **(A)** passive diffusion of nanomaterials through the intact ILM; **(B)** localized degradation of the ILM via proteases conjugated to silica microparticles; **(C)** facilitated diffusion of larger nanoparticles following ILM digestion. (Created in BioRender. Motyckova, L. (2026) https://BioRender.com/i3z5m8q)

## 2 Results and Discussions

We based our study on the porcine eye model, which shares extensive similarities with the physiology of the human eye^[30]^ and is potentially even closer to the human anatomy than most primates are.^[31]^ Our *ex vivo* experiments were conducted with several measures in place to preserve physiological relevance. First, we minimized the time elapsed *post-mortem*, initiating dissection within four hours after animal slaughter and completing all penetration experiments within six hours. Experiments performed over longer time durations sometimes led to visible signs of tissue degradation and drying. Second, we used a sclero-retinal tissue model in which a large area of the retina remained attached to the underlying sclera and was therefore less perturbed during the assessment of nanomaterial penetration. Notably, this approach prevented diffusion of nanoparticles around the ILM and their entry into the retina from the retinal pigment epithelium (RPE) side, which is inherently more permeable, thereby minimizing potential artifacts in the data. Third, we kept the retina hydrated at all times until the samples were frozen in liquid nitrogen, cryosectioned, fixed, and stained for visualization under a confocal microscope. The detailed preparation protocol is provided in the Experimental Section.

### 2.1 Size-dependent exclusion of nanomaterials by the ILM

We first evaluated the passive penetration of nanomaterials across the ILM by incubating retinas with differently-sized fluorescent probes (Fig. 2**A**). These included negatively-charged (carboxylated) nanoparticles made of polystyrene (50 nm, 100 nm, and 250 nm) or polymethacrylate (25 nm), and a fluorophore-labeled goat anti-rabbit IgG antibody (14 × 8 nm). No nanoparticle penetration beyond the retinal surface was detected, even for the smallest particles (25 nm) after 1 h of incubation (Fig. 2**C**; schematic in Fig. 2**B**). Prolonged *post-mortem* incubation (1 h, initiated 4 h after removal of nanoparticles), which would be expected to enhance tissue permeability, likewise did not permit nanoparticle passage through the ILM (Supplementary Fig. 1). In the examined retina sections, the fluorescence signal from the nanoparticles typically appeared as a uniform, sharp band approximately 12 µm thick, consistent across all particle sizes (25–250 nm). We reasoned that actual nanoparticle penetration through the ILM and diffusion in the tissue would occur in a more dispersed and irregular manner rather than the observed well-defined band and hypothesized that the observed fluorescence band is a consequence of the cryosectioning process. To investigate this, we sectioned the same sample at varying thicknesses (6, 12, and 30 µm). The results revealed a direct correlation between the band width and the section thickness, supporting the hypothesis that the observed fluorescence band corresponds to the thickness of the cryosectioned sample (Supplementary Fig. 2). We therefore conclude that carboxylated polystyrene and polymethacrylate particles do not cross the ILM unaided, but rather lie at the ILM interface.

**Figure 2:**
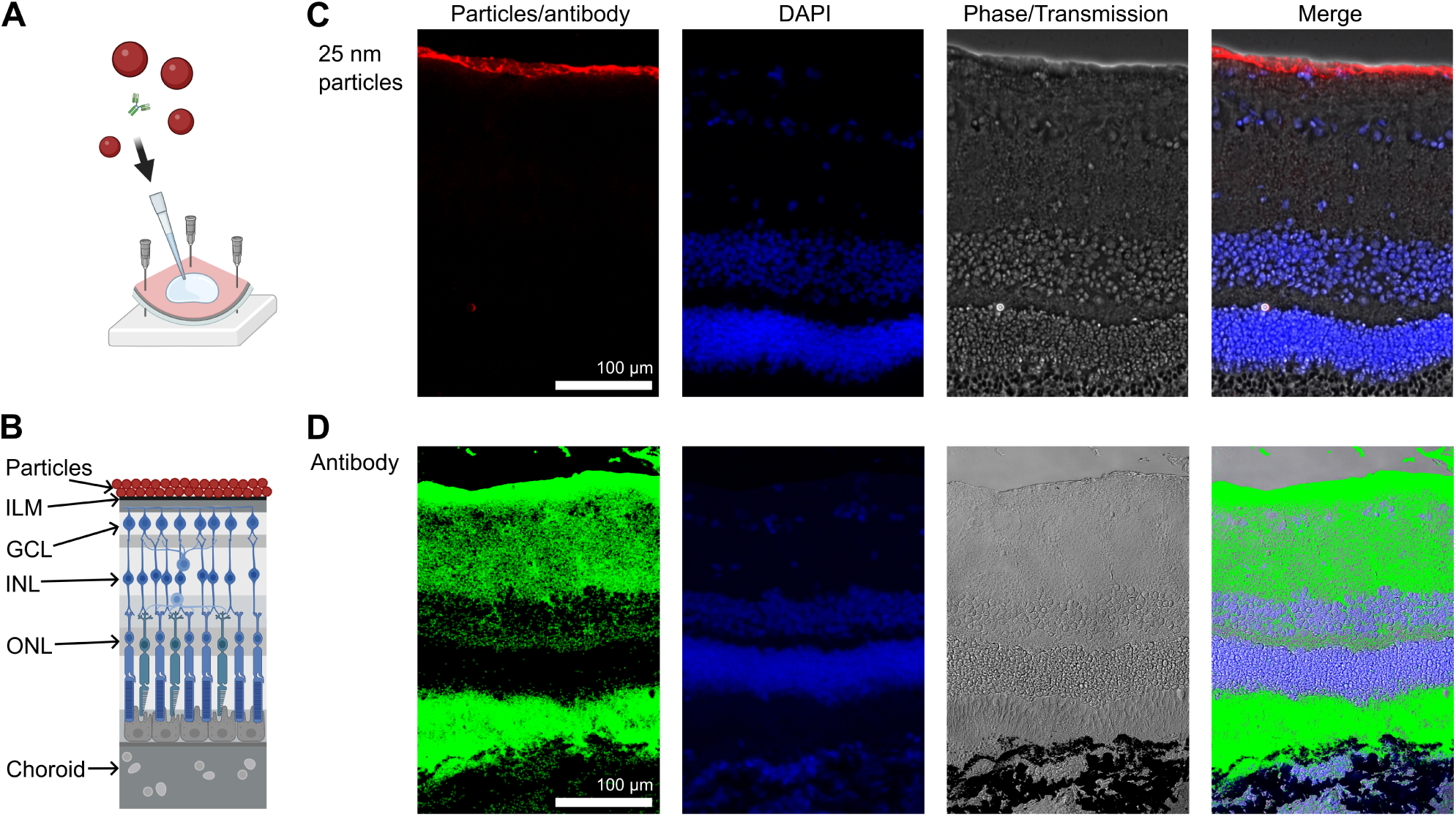
Assessment of ILM permeability to nanoscale carriers. **(A)** Schematic of a sclero-retinal explant used in permeability assays. **(B)** Schematic representation of retinal layers illustrating nanoparticles retained at the ILM without penetrating into retinal tissue. (Created in BioRender. Motyckova, L. (2026) https://BioRender.com/1njthbs) **(C)** Representative fluorescence images showing retention of 25 nm fluorescent carboxylated polymethacrylate nanoparticles at the ILM. **(D)** In contrast, fluorophore-labeled goat anti-rabbit IgG secondary antibody (14 nm × 8 nm) successfully crossed the ILM and diffused into the retina. Pixels with antibody-associated signal (ratio intensities > 1) are highlighted with green outlines. Columns display (from left to right): red fluorescence (nanoparticles) or highlighted antibody signal (green), DAPI staining, phase/trasmission images, and merged images. Abbreviations: GCL – Ganglion Cell Layer, ILM - Inner Limiting Membrane, INL – Inner Nuclear Layer, ONL – Outer Nuclear Layer.

In contrast, the fluorophore-labeled antibody (14 nm × 8 nm) successfully traversed the ILM and diffused through the retinal layers, reaching the RPE within 1 h of incubation, as confirmed by confocal microscopy imaging (Fig. 2**D**). To ensure accurate detection of penetration, we implemented an imaging strategy designed to minimize interference from tissue autofluorescence. This was necessary as the observed autofluorescent signal varied significantly across, and even locally within, samples, potentially masking the weaker fluorescence signals from the probes. During image acquisition with the same excitation wavelength, an additional detection channel was recorded simultaneously, capturing emissions at longer wavelengths than those of the fluorophore of interest (see Experimental Section for details). Given that autofluorescent background signals are largely independent of detection wavelength, the ratio of signal intensities between the fluorophore channel and the longer-wavelength channel was calculated on a per-pixel basis. Pixels with ratio values greater than one were considered to represent true fluorophore-associated signal, thereby isolating specific nanomaterial fluorescence from background noise. For enhanced visualization, these pixels are highlighted with green outlines in Fig. 2**D**.

Overall, these findings suggest that the ILM imposes a size exclusion threshold for negatively charged particles between ≈ 10 and 25 nm. Given that the smallest transfection agents used in retinal gene therapy are larger than this threshold, passive diffusion alone is unlikely to be sufficient for effective delivery. This underscores the need for advanced targeted strategies to facilitate transfection agent transport across the ILM after intravitreous administration of gene therapy agents. In this context, active strategies that facilitate the transport of nanoscale particles across the ILM are of interest. One approach is to utilize external forces acting on particles through magnetic^[32]^ or electric fields^[33]^, while others aim to transiently disrupt the ILM via physical^[34,35]^ or biochemical^[27,36,37]^ means. Notably, Peynshaert *et al.* developed a light-based method in which indocyanine green enabled laser-mediated ILM ablation, later refined through nanoparticle encapsulation for more localized disruption.^[34,35]^

In this study, we focus on localized enzymatic degradation as a general approach to increase the permeability across the ILM without the need for specialized equipment.

### 2.2 Localized ILM degradation by collagenase immobilized on microparticles

For the localized enzymatic degradation, we selected collagenase type IV, because it selectively degrades type IV collagen^[38]^, the principal structural component of the ILM.^[9,10]^ Previous studies have shown that proteolytic enzymes can enhance ILM permeability^[27,36,37]^, but these approaches typically required high enzyme concentrations that can compromise the structural integrity of the underlying retina, are toxic to retinal cells and may cause hemorrhage due to degradation of blood vessel basal membranes. To address these limitations, collagenase IV immobilized on polydopamine nanoparticles has recently been shown to induce vitreous liquefaction while preserving the retina’s structure^[39]^. In this work, we employed an alternative immobilization strategy, coupling collagenase IV to silica microparticles with the goal of locally degrading the ILM to enhance nanoparticle transport, while maintaining the structural integrity of the retina.

Enzyme immobilization was achieved through a multi-step surface functionalization protocol, adapted from Marcelino *et al.*.^[40]^ First, SiO_2_ microparticles (Fig. 3**A**) were silanized with an APTES-ATRP initiator, providing active sites for polymer growth. Next, pDEAEMA polymer brushes were grafted from the bead surface, creating a functional scaffold for enzyme conjugation (Fig. 3**B**, **D**). Finally, collagenase IV was immobilized on the SiO_2_-Polymer particles through incubation at pH 6.4, which promoted electrostatic interactions between the positively charged pDEAEMA and negatively-charged collagenase IV (Fig. 3**C**) (see Methods for further experimental details). ζ-potential analysis confirmed the stepwise charge evolution: bare SiO_2_ microparticles were negatively charged, polymer grafting shifted the surface potential to positive values, and subsequent enzyme immobilization restored a net negative charge (Fig. 3**E**). Notably, this final negative ζ-potential of the SiO_2_@Polymer–Enzyme particles is consistent with prior reports and deliberately chosen, based on the findings that negatively charged carriers exhibit enhanced penetration into the retina.^[24–26]^ The activity of immobilized enzyme on the surface of SiO_2_ particles was determined using a collagen IV activity assay to 0.013 U for 1 mg of particles, which correlates to ≈0.82 µg of free enzyme.

**Figure 3:**
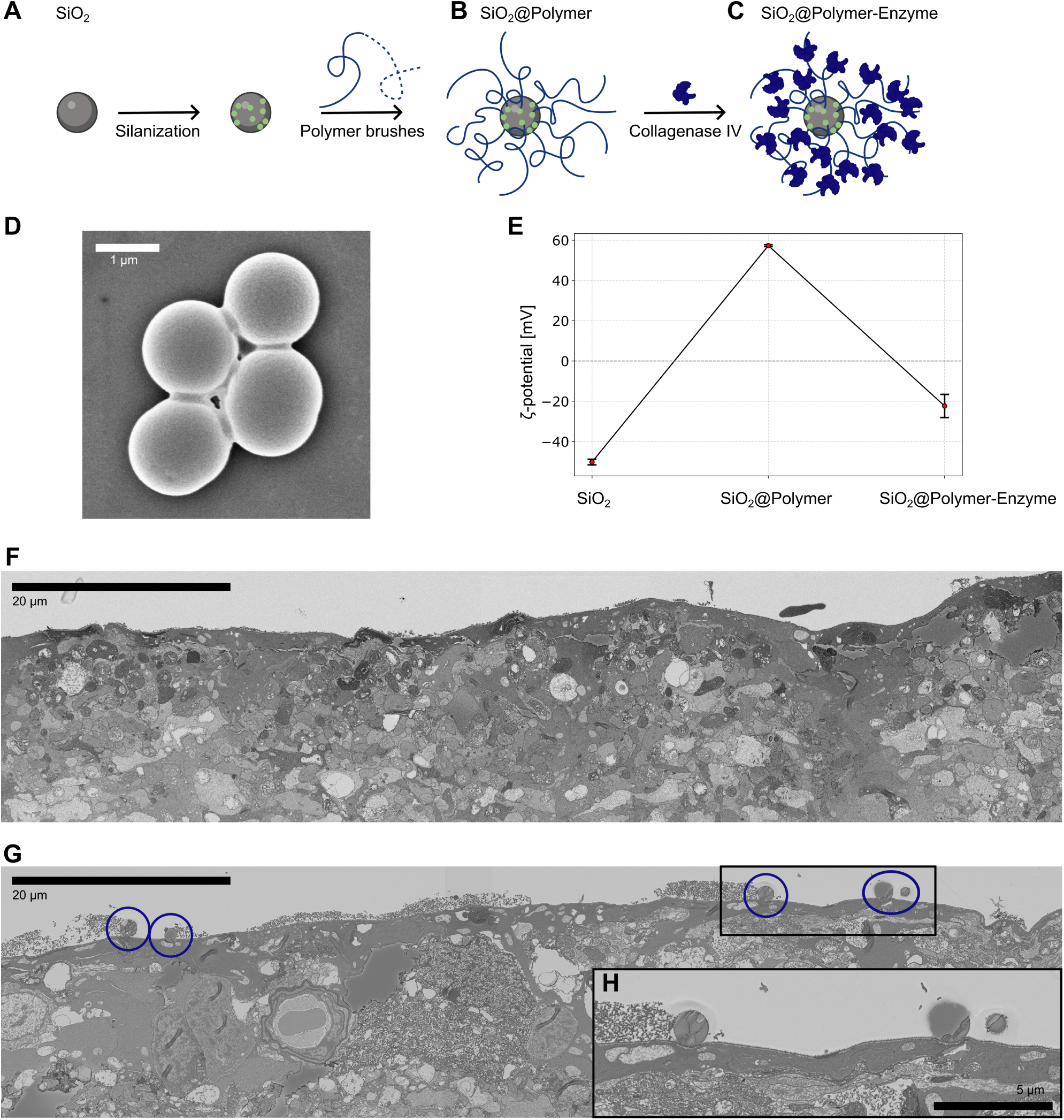
Immobilization of collagenase IV on silica microparticles. Schematics: **(A)** SiO_2_ particles, their silanization with an APTES-ATRP initiator, and subsequent surface grafting of pDEAEMA polymer brushes; **(B)** SiO_2_@Polymer particles and subsequent immobilization of collagenase IV onto the polymer scaffold; **(C)** SiO_2_@Polymer–Enzyme particles. (Created in BioRender. Motyckova, L. (2026) https://BioRender.com/7ea9beq) Characterization: **(D)** SEM image of SiO_2_@Polymer particles; **(E)** ζ-potential measurements of SiO_2_, SiO_2_@Polymer, and SiO_2_@Polymer-Enzyme particles; **(F)** electron microscopy of retinal tissue after application of non-functionalized SiO_2_@Polymer particles; **(G)** retinal tissue following treatment with SiO_2_@Polymer-Enzyme particles (microparticles are highlighted with blue outlines); and **(H)** magnified view of panel (G).

Next, the SiO_2_@Polymer and SiO_2_@Polymer-Enzyme particles were incubated with scleroretinal explants for 1 h (see Methods for details), and the retinal interface was examined using ultra structural electron microscopy (Fig 3**F**–**H**). In explants exposed to SiO_2_@Polymer-Enzyme particles, multiple microparticles were consistently detected at the ILM (≈ 7 microparticles per 100 µm of ILM, Fig. 3**G**–**H**), whereas no microparticles were detected at the retinal surface in control samples lacking collagenase (Fig. 3**F**). Notably, microparticles were absent within the retinal layers under both conditions. The selective retention of SiO_2_@Polymer-Enzyme particles at the ILM provides an indication of localized enzymatic activity.

To further investigate whether SiO_2_@Polymer-Enzyme particles induce localized degradation of the ILM, we performed a two-step incubation experiment. Sclero-retinal explants were first exposed to SiO_2_@Polymer-Enzyme particles, followed by incubation with 25 nm fluorescent carboxylated polymethacrylate particles, which served as markers to detect ILM disruption. This treatment indeed confirmed localized ILM degradation and enabled trans-ILM delivery of the nanoparticles (Fig. 4 and Methods for details). Nanoparticle penetration was assessed using the two-channel imaging and pixel ratio analysis described earlier, with pixels exhibiting ratio values greater than one classified as fluorophore-associated signal and highlighted with yellow outlines in Fig. 4. In line with our previous experiments (Fig. 2**C**), in the buffer-only control, where sclero-retinal explants were incubated in HBSS supplemented with Ca^2+^ prior to nanoparticle application, no nanoparticle penetration beyond the ILM was observed (Fig.4**A**). Importantly, no significant increase in ILM permeability to nanoparticles was observed when explants were incubated with SiO_2_@Polymer particles (Fig.4**B**). Instead we observed an interaction of polymer brushes with the nanoparticles, leading to co-localization of both when no enzyme was present, as a result of the electrostatic interactions between oppositely charged particles. In contrast, incubation with free collagenase IV solution (Fig.4**C**) or collagenase-immobilized microparticles (Fig.4**D**) enabled subsequent robust nanoparticle penetration into the retinal layers, demonstrating that enzymatic activity of collagenase IV was required for effective ILM degradation and subsequent nanoparticle penetration.

**Figure 4:**
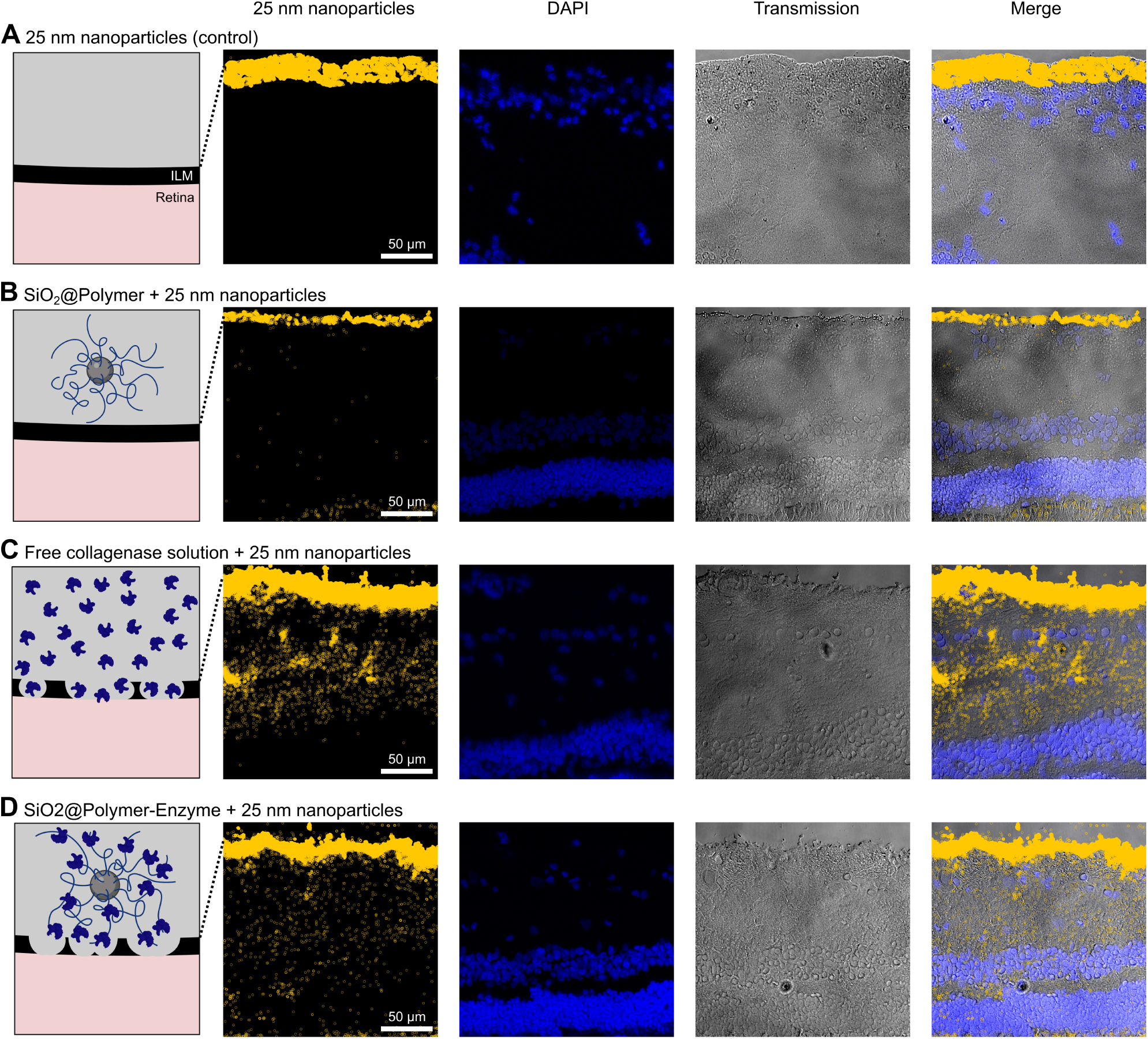
Collagenase IV treatment facilitates retinal penetration of 25 nm polymethacrylate nanoparticles. Representative data from sclero-retinal explants are shown for four experimental conditions: (A) nanoparticles alone, (B) polymer-functionalized silica microparticles without enzyme followed by nanoparticle application, (C) free collagenase IV solution followed by nanoparticle application, and (D) polymer-functionalized silica microparticles carrying immobilized collagenase IV followed by nanoparticle application. For each condition, the first column shows a schematic of the treatment prior to nanoparticle application, followed by confocal images displaying pixels with ratio intensities > 1 (fluorophore-associated nanoparticle signal, highlighted with yellow outlines), DAPI staining, transmitted light images, and merged images. The scale bars apply to all images within the same row. (Created in BioRender. Motyckova, L. (2026) https://BioRender.com/crpjbp1)

To complement these qualitative observations, we performed a quantitative analysis of nanoparticle penetration across different experimental conditions. Specifically, the proportion of pixels classified as fluorophore-associated signal (ratio > 1) was calculated across multiple confocal images obtained from three independent explants for each condition (Table 1). The experimental groups comprised four treatments in which the indicated additives were diluted in HBSS buffer, followed by 25 nm fluorescent nanoparticle application: (A) control (no additives), (B) SiO_2_@Polymer, (C) free collagenase IV solution, and (D) SiO_2_@Polymer–Enzyme. For each explant, the mean number of fluorophore-associated pixels was calculated from confocal images cropped in a distance of 13.5 µm beneath the ILM. The average for the three independent explants per condition are shown in Table 1. Exposure to free collagenase IV significantly increased subsequent nanoparticle penetration into the retinal layers (mean 678 pixels) where nanoparticles alone (mean 67 pixels) show almost no penetration at all. Similarly, treatment with SiO_2_@Polymer-Enzyme particles significantly increased nanoparticle diffusion (mean 377 pixels) relative to both nanoparticles alone and polymer-functionalized particles without enzyme (mean 120 pixels). These results confirm that enzymatic activity, whether in solution or immobilized on microparticles, facilitates trans-ILM transport of 25 nm nanoparticles. The increase in particle penetration in the case of SiO_2_@Polymer-Enzyme to comparable levels with free enzyme solution is remarkable, considering the 1000-fold lower utilized activity. This substantial decrease in effective enzyme activity should reduce toxic off target effects, supported by the restricted movement of the immobilized enzymes preventing degradation in deeper located tissues.

**Table 1:** Quantitative analysis of 25 nm particle penetration across different experimental treatments prior to nanoparticle application. Pixel counts greater than 1 (“Pix.” in the table) were extracted from selected cross-sections of multiple images (“Image No.” in the table) for three explants per condition. The mean pixel values across the three explants are reported as “Average Pix.” for each treatment.

| Condition | Explant 1<br>Pix. (Image No.) | Explant 2<br>Pix. (Image No.) | Explant 3<br>Pix. (Image No.) | Average<br>Pix. |
| --- | --- | --- | --- | --- |
| Control | 2(6) | 91(11) | 109(13) | 67 |
| SiO<sub>2</sub>@Polymer | 34(15) | 322(12) | 5(13) | 120 |
| Free collagenase solution | 1042(3) | 120(15) | 871(15) | 678 |
| SiO<sub>2</sub>@Polymer-Enzyme | 903(23) | 92(8) | 138(9) | 377 |

### 2.3 Evaluation of biocompatibility in an *in vivo* porcine model

Considering the prospective future application of collagenase-decorated microcarriers for localized ILM degradation *in vivo*, we evaluated the biocompatibility of SiO_2_@Polymer-Enzyme in a relevant *in vivo* porcine model. As part of a larger study (Ethical Approval no. 35-9185.81/G-11/24 and 35-9185.81/G-156/25, Regierungspräsidium Karlsruhe), the eyes of two German Landrace pigs were monitored following intravitreal injections. One animal received SiO_2_@Polymer-Enzyme in one eye, and the other animal received SiO_2_@Polymer in one eye, with the respective contralateral second eyes serving as controls. Eyes were injected with 100 µL of suspension containing approximately 5.6 × 10^8^ particles, fabricated identically to those used in the *ex vivo* experiments and following procedures approved by the local ethics committee (see Supplementary Note 1 for details). Biocompatibility assessments were conducted at baseline (prior to injection) and at two-week intervals up to six weeks post-injection. Comprehensive ophthalmic evaluations—including fundus photography, optical coherence tomography (OCT), full-field electroretinography (ff-ERG), and intraocular pressure (IOP) measurements—were performed in the anesthetized animals. Both animals completed the full six-week study period without systemic adverse effects related to the injected substances.

Throughout the six-week observation period, no morphological changes regarding the posterior eye segment and no signs of inflammation, atrophy or hemorrhage occurred in the animals, see Figure 5 & Supporting Figure 4 and Supporting Note 1. Similarly, the OCT scans of the two treated animals revealed no alterations in retinal ultrastructure over the course of the study (Figure 5).

**Figure 5:**
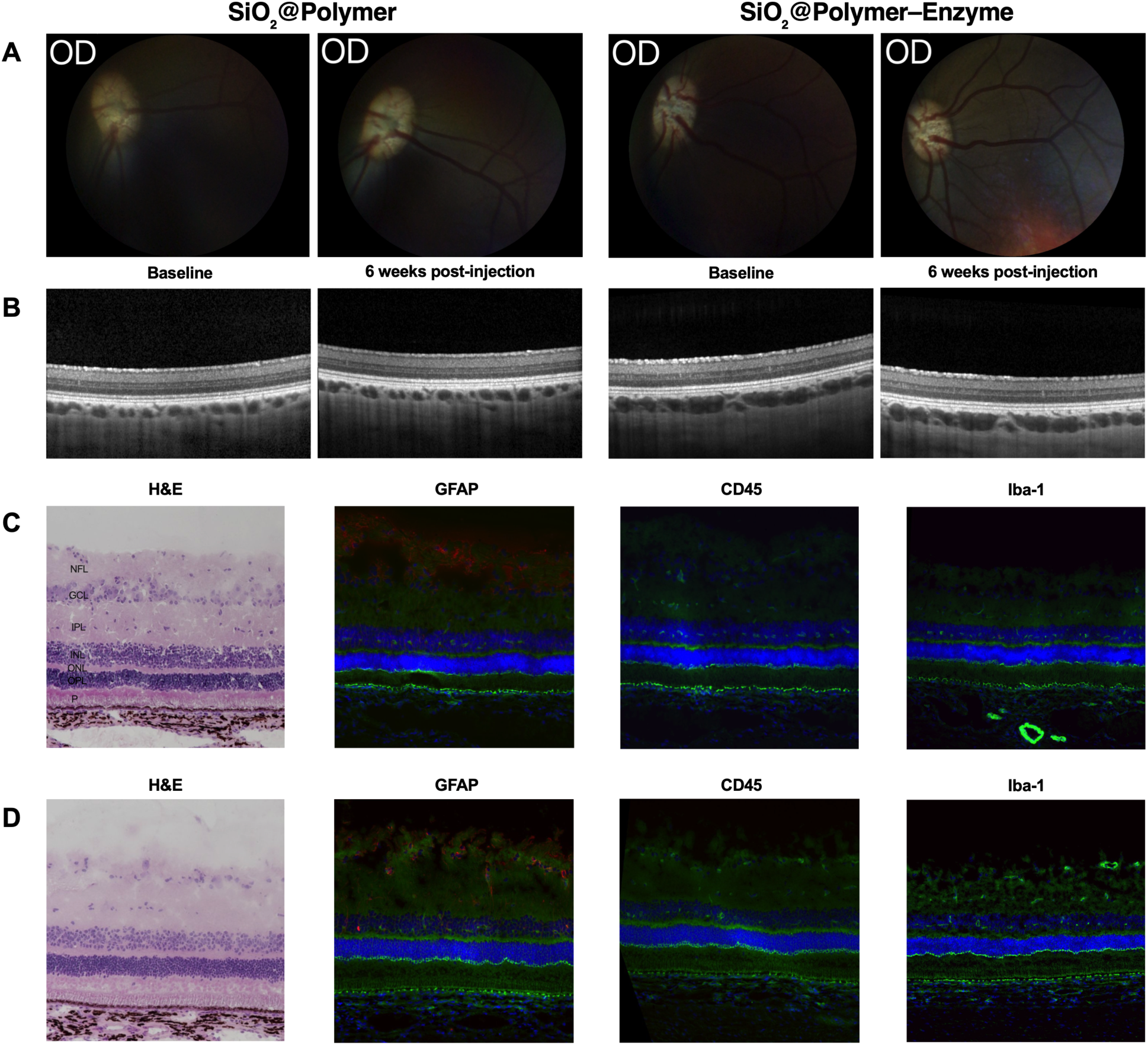
Biocompatibility of SiO_2_@Polymer-Enzyme and control SiO_2_@Polymer studied in an *in vivo* pig model over the course of six weeks. **(A)** Fundus photographs: treated right eyes showed no morphologic alterations of the fundus of both animals after six weeks compared to baseline indicating no significant retinal toxicity. **(B)** Representative OCT sections: the treated right eye of both animals after six weeks compared to baseline revealed no changes in retinal ultrastructure and no inflammatory or neovascular reaction. **(C-D)** H&E and immunohistochemical staining of the eyes treated with SiO_2_@Polymer **(C)** and with SiO_2_@Polymer-Enzyme **(D)**: no structural alterations in the treated eyes were detected in H&E staining and no signs of glial proliferation were indicated by GFAP-staining **(red)**. Microglial or leucocyte infiltration into retinal structures did not occur according to Iba-1 **(red)** and CD-45 staining **(red).** Abbreviations: GCL – Ganglion Cell Layer, INL – Inner Nuclear Layer, IPL – Inner Plexiform Layer, NFL – nerve fiber layer, ONL – Outer Nuclear Layer, OPL – Outer Plexiform Layer, P – Photoreceptor layers.

Following completion of the six-week study, animals were euthanized, and the enucleated eyes were processed for histology and immunohistochemistry. Results from Hematoxylin-Eosin-staining (H&E) showed no signs of atrophy, loss of cell density or other structural alterations in the retina (Fig. 5**C**, **D**). Furthermore, no significant invasion of leucocytes (CD45), microglial cell activation and migration (ionized calcium-binding adapter molecule 1, Iba-1) or activation of Müller glia (GFAP staining) were apparent. (Fig. 5**C**, **D**).

Based on these findings, we conclude that the SiO_2_@Polymer and SiO_2_@Polymer-Enzyme particles used for the penetration studies exhibit excellent *in vivo* biocompatibility, supporting their suitability as potential components of targeted ocular drug delivery systems.

## 3 Conclusion

In this study, we first addressed the size threshold for nanoparticles to penetrate the inner limiting membrane of the eye. We also demonstrated how enzymatically-active microcolloids can be used to facilitate penentration of nanoparticles by locally degrading the ILM in a porcine model. In particular, our results indicate that the major diffusion barrier, the ILM, has a pore size below 25 nm, substantially smaller than the dimensions of most currently engineered drug delivery platforms. This discrepancy between available technologies and biological constraints highlights the need for fundamental studies using appropriate model systems to generate reliable and translatable data. The stringent size limitation poses a significant challenge, particularly for delivery of larger biomolecules such as DNA, and motivates the exploration of alternative strategies to overcome passive diffusion constraints. Here, we investigated localized enzymatic degradation of the ILM as a means to increase retinal permeability. Based on our *ex vivo* model, we demonstrate that silica microparticles functionalized with immobilized collagenase IV can facilitate the penetration of 25 nm negative particles into all retinal layers from the vitreous side. Future studies should explore longer exposure times, varying doses, and ultimately *in vivo* validation to assess the translational potential of this approach. While previous work has shown that high doses of proteases can adversely affect the underlying retinal nerve fibers—a finding consistent with our observations—our *in vivo* experiments reveal no detrimental impact on retinal function, morphology, or topology following intravitreal injection of polymer-decorated silica microparticles, with or without immobilized collagenase IV. These results suggest that enzymatic functionalization of nanoparticles represents a promising strategy for minimally invasive transport across the vitreoretinal interface. We expect that the use of enzymatically-active microparticles presents a general strategy to aid the transport of nanotherapeutics and gene carriers across biological barriers.

## 4 Experimental Section

### 4.1 Materials

Silica microparticles with diameter of 1.5 µm (sicastar^®^) were acquired from micromod Partikeltechnologie. Similarly, fluorescent carboxylated polystyrene/polymethacrylate particles (micromer^®^-redF) were purchased from micromod Partikeltechnologie in a range of sizes: polymethacrylate (25 nm) and polystyrene (50 nm, 100 nm, 250 nm). Goat anti-Rabbit IgG secondary antibody conjugated to Alexa Fluor 488 and 4’,6-diamidino-2-phenylindole (DAPI) fluorescent dye was obtained from Invitrogen. Hank’s balanced salt solution (HBSS) with calcium and magnesium was acquired from Gibco™ (Cat. No. 24020117). Collagenase type IV (C4-BIOC), (+)-Sodium L-ascorbate, α-Bromoisobutyryl bromide (BIBB), tris[2-(dimethylamino)ethyl]amine (Me_6_TREN), 2-(Diethylamino)ethyl methacrylate (DEAEMA), APTES were purchased from Sigma-Aldrich. Tissue-Tek^®^ O.C.T. Compound was acquired from Sakura Finetek USA, Inc. SuperFrost^®^+ cover glasses were purchased from R. Langenbrinck GmbH. FluorSave Reagens was purchased from Millipore. Reagents for electron microscopy were bought from Sigma (Pipes, tannic acid, potassium ferricyanide), Science Services (glutaraldehyde, paraformaldehyde, OsO4, uranylacetate, lead citrate), and Serva (glycid ether, DDSA, MNA, BDMA).

### 4.2 Eye dissection

Fresh porcine eyes were obtained from a local slaughterhouse and maintained at 4°C following enucleation until dissection, which was performed within four hours *post-mortem*. Upon removal of all extraocular tissue, the eyes were disinfected in 70% ethanol and rinsed with sterile phosphate-buffered saline (PBS). The sclera was punctured with a surgical scalpel (Supplementary Fig. 3**A**), creating an incision that allowed the insertion of surgical scissors for the excision of the anterior segment of the eye (Supplementary Fig. 3**B**). The vitreous humor was carefully extracted by gently tilting the eye cup and delicately pulling the vitreous using tweezers (Supplementary Fig. 3**C**). The posterior eye cup was subsequently bisected, ensuring that the optic nerve remained on one side of the cut (Supplementary Fig. 3**D**), yielding two *ex vivo* sclero-retinal samples per eye (Supplementary Fig. 3**E**). Each sample was carefully affixed to a styrofoam plate using needles to prevent collapse of the tissue structure (Supplementary Fig. 3**F**). Nanomaterials or microparticles with or without enzymes, suspended in Hank’s balanced salt solution (HBSS) supplemented with Ca^2+^, were carefully applied to the sample using a pipette (Supplementary Fig. 3**G**). The samples were then incubated at 37*^◦^*C (Supplementary Fig. 3**H**). Following incubation, a central region of each sample was excised using surgical scissors (Supplementary Fig. 3**I**) and embedded in Tissue-Tek O.C.T. medium within a mold constructed from aluminum foil (Supplementary Fig. 3**J**). To ensure uniform freezing, the molds were placed on an aluminum foil raft floating on liquid nitrogen (Supplementary Fig. 3**K**). Once frozen, the samples were stored at −80°C until cryosectioning.

### 4.3 Silica particle polymer brush functionalization

The polymer brushes were grafted from the silica particle surface according to the literature.^[40]^ The surface of silica particles was modified with an APTES-ATRP (atom transfer radical polymerization) initiator before performing a pDEAEMA brush polymerization. Silica particles (50 mg) were re-dispersed in 15 ml of ethanol. After addition of ATPES (1 ml) the dispersion was heated under reflux for 1 h and subsequently washed with an ethanol/toluene mixture with gradually increasing amounts of toluene (2:1, 1:1, 1:2). The particles were then re-dispersed in dry toluene and transferred to a clean flask. After addition of triethylamine (200 µl) and BIBB (150 µl) the dispersion was stirred overnight in a chemical hood with a septum with a needle inserted to allow gas exchange. The APTES-ATRP functionalized particles were washed in toluene/ethanol mixture with gradually increasing amounts of ethanol (3:1, 2:1, 1:1, 1:2, 1:3) and re-dispersed in 1 ml of ethanol.

For the polymerization, the APTES-ATRP functionalized particles (10 mg, 200 µl) were combined with CuSO_4_ (400 µl, 0.1 mg · ml*^−^*^1^ in water), sodium ascorbate (200 µl, 0.5 mg · ml*^−^*^1^ in water), Me_6_TREN (16.7 µl) and DEAEMA (109.2 µl, 0.7 M, in ethanol) and rotated for 1 h. Subsequently the particle aggregates were transferred to water through dialysis (MWCO 3500 Da) for 1 day with repeated water exchange. The particle aggregates were then washed in phosphate buffer at pH 1 and sonicated for 30 min. Finally the particles were re-dispersed in phosphate buffer at pH 6.4. Enzyme (100 µg per mg) was immobilized on the particles by incubation for 12 h/ON at 4 °C and subsequent washing (4x).

The phosphate buffer consisted of 2.5 mM Na_2_HPO_4_ and 50 mM NaCl. The pH was adjusted using diluted HCl (aq).

### 4.4 Zeta potential

Zeta potential measurements were performed on a Malvern Zetasizer Nano using DTS1070 cuvettes. Particles were diluted in water to appropriate concentration and measured in triplicate.

### 4.5 Enzyme activity

Enzyme activity was quantified using EnzChek Collagenase Assay Kit with DQ collagen type IV from human placenta, fluorescein conjugate (Invitrogen) according to the manufacturer’s manual. Fluorescence was measured using a plate reader at 37 °C with shaking before each measurement. All values were corrected using a negative control without enzyme. Only the initial, linear slope of the graphs was considered in the analysis.

The activity of polymer-functionalized silica microparticles carrying immobilized collagenase IV was measured to 0.13 U/mg particles, which corresponds to the activity of ≈0.82 µg of free enzyme when assuming a moderate enzyme activity of 160 Mandl-U/mg.

### 4.6 Incubation of sclero-retinal explants

Sclero-retinal explants were incubated for 1 h at 37 *^◦^*C in HBSS supplemented with 1.26 mM Ca^2+^ and exposed to different conditions, followed by a 1 h incubation with 25 nm fluorescent carboxylated polymethacrylate nanoparticles under the same conditions. Calcium was included as a cofactor for enzyme activity, and the collagenase IV blend with clostripain used exhibits 125–250 Mandl-U/mg (manufacturer specification). Four experimental groups were tested: (i) collagenase IV solution (5 mg, 800 Mandl − U per explant, explants washed with HBSS prior to nanoparticle exposure), (ii) HBSS with Ca^2+^ (buffer control), (iii) polymer-functionalized silica microparticles carrying immobilized collagenase IV (1 µg, 0.13 Mandl − U per explant), and (iv) polymer-functionalized silica microparticles without enzyme (bead control).

### 4.7 Histological processing of sclero-retinal explants

The molds containing the sclero-retinal explants were sectioned into 12 µm slices using a CryoStar NX50 cryotome (Thermo Scientific) and subsequently transferred onto a coverslip. The sections were fixed by applying methanol for 10 min. The slides were then washed in Tris-buffered saline (TBS) for a minimum of 10 min before staining with DAPI for 5 min. After two washes in TBS and one wash in water, 10 min each, the samples were dried for 15 min. The samples were mounted using FluorSave reagent.

### 4.8 Quantitative analysis of nanoparticle penetration

For each experimental condition, three independent sclero-retinal samples were obtained from separate porcine eyes on at least two different days. The cryosections were imaged with a confocal microscope (LSM 880, Zeiss) using a a Plan-Apochromat 20×/0.8 NA air objective with 405 nm and 561 nm lasers in two separate channels. Signal collection for ratio imaging (561 nm laser) was set for following ranges: Ch1: 569-631 nm, Ch2: 668-727 nm, Ch3: T-PMT. The acquisition settings for Ch1 and Ch2 were otherwise identical.

Pixel-wise ratios (Ch1/Ch2) were calculated to distinguish the particle fluorescence (Ch1) and correct for prominent and inconsistent autofluorescence background (Ch2) signals from the ocular tissue. The ratio Ch1/Ch2 was used to normalize the background signal to values ≤ 1, which could be subtracted.

From these ratio images, a 500 × 500 pixel (104 x 104 µm) region was cropped, starting approximately 65 pixel (13.5 µm) beneath the retinal surface to exclude the superficial, overlapping particle band along the ILM. Pixels with a ratio exceeding 1 were classified as fluorophore-associated signal and quantified across the cropped region. For each sample, multiple confocal images (ranging from 3 to 26) were analyzed. Outlier images were identified and excluded using Grubbs’ statistical test, resulting in 3–23 images per sample included in the final analysis. The mean number of fluorophore-associated pixels was then calculated for each sample. Finally, the average number of pixels with ratio > 1 was computed for each experimental condition based on three independent samples.

### 4.9 Explant fixation and embedding for scanning electron microscopy (SEM)

After one hour treatment, samples were fixed with 1.25 % glutaraldehyde and 2 % paraformaldehyde in 0.1 M PIPES buffer, pH 6.9 and left overnight. After washing with PIPES buffer, samples were incubated in 1 % tannic acid in 0.1 M PIPES buffer for 1 hour, followed by heavy metal staining with 1 % OsO_4_ and 1.6 % potassium ferricyanide (K_3_[Fe(CN)_6_]) in 0.1 M PIPES buffer for 1 hour on ice. For the samples incubated with glass microparticle concentrations of tannic acid and OsO_4_ were 0.5 %. All samples were block-stained overnight in 2 % uranyl acetate in 25 % ethanol. Dehydration with ethanol (50 %, 70 %, 90 %, and 100 %) for 15 min each was followed by 15-minute incubations in a 1:1 ethanol-acetone mixture and 100 % dried acetone. After infiltration with 30 % and 70 % Epon in acetone for 2 h each, samples were left in 100 % Epon overnight, then embedded in molds with fresh Epon and polymerized at 60 °C for 2 days. Epon was prepared by mixing 21.2 g glycid ether 100 with 14.8 g dodecenyl succinic anhydride (DDSA), 9.2 g methyl nadic anhydride (MNA) and 1.2 g benzyldimethylamine (BDMA) as accelerator.

### 4.10 Sectioning and scanning electron microscopy (SEM)

Sample blocks were trimmed and sectioned with diamond knives (DiATOME, Switzerland) using an ultramicrotome (PowerTome PT-PC, RMC Boeckler, US). Individual sections were on average 100 nm thick. Sections were placed on pieces of silicon wafers (2 × 2 cm) and post-stained with 3 % lead citrate for 10 min followed by 3 % uranyl acetate for 5 min. SEM imaging was performed using a ULTRA 55 with software packages SmartSEM and Atlas 5 (Carl ZEISS Microscopy, Germany). Imaging parameters were 1.5 keV primary energy, 20 µm aperture, with SE2, EsB, and InLens detectors.

### 4.11 Biocompatibility examination in an *in vivo* porcine model

The *in vivo* study was carried out at the Interfaculty Biomedical Research Facility (IBF) of Heidelberg University.

Sedation of the animals for the examinations was performed with azaperon (Sedanol, WDT, Garbsen, Germany) (6 mg/kg) applied intramuscularly. Ketamine (CP Pharma, Burgdorf, Germany) (11 mg/kg) and midazolam (Hameln Pharma, Hameln, Germany) (2 mg/kg) administered intramuscularly was used to induce anesthesia. For maintaining the anesthesia, Isoflurane (CP Pharma, Burgdorf, Germany) 2 % (v/v) was used. Medical mydriasis for the biocompatibility examinations was induced using cyclopentolate (0,5 %), epinephrine (5 %) and tropicamide (1 %) as needed.

OCT was performed using the Spectralis Flex OCT system (Heidelberg Engineering GmbH, Heidelberg, Germany). Scans were located in visual streak, the cone-enriched center of the porcine retina corresponding to the human macula. At the 6 weeks timepoint, angiographic examination with intravenous administred fluorescein (10%, Alcon, Freiburg, Germany) and indocyanine green (Verdye 5 mg/mL, Mediconsult, Roggwil, Switzerland) was performed using the Spectralis Flex system. Images of the peripapillary region and the visual streak were acquired about 15 min after intravenous administration of the contrast agents. For fundus photography, the ClearView2 veterinary fundus camera (Optibrand Ltd., Fort Collins, USA) was used and IOP measurement were conducted with the iCare Tonovet Plus (iCare Finland Oy, Vantaa, Finland).

For ERG measurements, the RETevetTM (LKC Technologies, Gaithersburg, USA) system, was used following the Dog, Cat, Nonhuman Primate ISCEV 6 Step testing procedure. The active electrode was placed on the cornea using lubricating Carbomer (Vidisic®, Dr. Gerhard Mann chem.-pharm. Fabrik GmbH, Berlin, Germany) for corneal protection. The reference electrode was placed temporal near the lateral canthus and the grounding electrode was positioned on the pig’s forehead. Both eyes were dark adapted for 15 minutes. The testing protocol is visualized Supplementary Table 1.

### 4.12 Histological investigation of *in vivo*–treated porcine models

*Post-mortem*, injected eyes as well as left control eyes were enucleated. The vitreous cavity was filled with pre-cooled isopentane (>99%, Carl-Roth) and Tissue Freezing Medium (Tissue Freezing Medium, LEICA Biosystems) was used to fill the posterior eyecup before freezing the sample in liquid nitrogen. Cutting of the frozen eyes into 5-7 µm thick slices was performed by cryostat (Leica CM1850, Germany) cooled down to −16 to −19°C. Slices were mounted on Superfrost Plus microscopic slides (Epredia, Germany) and air-dried. Prior to Hematoxylin-Eosin-staining (H&E), slides were rinsed in distilled water. After immersion in hematoxylin solution (acidic Mayeŕs hematoxylin solution, Carl-Roth GmbH + Co. KG, Karlsruhe, Germany) for six minutes, slides were blued in running tap water for 15 minutes and then rinsed in distilled water for two minutes. Counterstaining was performed with eosin G solution (0.5%, Carl-Roth GmbH + Co. KG, Karlsruhe, Germany), supplemented with one drop of glacial acetic acid (99.8 – 100%, Bernd Kraft GmbH, Den Haag, Netherlands) for three minutes. Afterwards, slides were rinsed with tap water, dehydrated in ascending ethanol solutions (70%, 96% and 100%; ZENTRALBEREICH Neuenheimer Feld, Heidelberg, Germany) and cleared in ROTIHistol (Carl-Roth GmbH + Co. KG, Karlsruhe, Germany). For coverslipping, ROTIHistokitt (Carl-Roth GmbH + Co. KG, Karlsruhe, Germany) and glass coverslips (24 x 50 mm, borosilicate glass, VWR International, Radnor, USA) were applied. Fixation of Immunohistology slides was performed using Acetone (>99,5%, Carl-Roth GmbH + Co. KG, Karlsruhe, Germany) at −10°C. After air-drying, each section was circled using a hydrophobic barrier pen (Dako® PAP Pen, Aglient Technologies, Santa Clara, USA). Blocking was performed by application of 10% horse serum (S-2000, Vector Laboratories, Burlingame, USA), diluted in 1% BSA/PBS for 30 mins in a dark humid chamber. Blocking solution was removed by tapping the slides onto a tissue. Primary antibodies were diluted in 1% PBS (1:100). Anti-CD45 (clones 2B11 + PD7/26, Dako, Agilent Technologies, Santa Clara, USA), anti-GFAP (clone GF 12.24, PROGEN, Heidelberg, Germany), anti-IBA1 (HPA049234, Atlas Antibodies, Sigma-Aldrich, USA) and one control vial with PBS was applied, each on one slice. The slides were stored at 4°C overnight. After staining, slides were washed twice in 1% PBS for five minutes and Cy3-conjugated donkey anti-mouse secondary antibody (Jackson ImmunoResearch Laboratories, West Grove, USA) for GFAP and CD45 primary antibodies, Cy3-conjugated donkey anti-rabbit secondary antibody (Jackson ImmunoResearch Laboratories, West Grove, USA) for IBA1 primary antibody, DAPI (D9542, Sigma-Aldrich, Merck, Germany) and Phalloidin (CytoPainter, ab176753, Abcam Cambridge, UK) were applied all together diluted in concentrations of 1:200, 1:200, 1:1000 and 1:200 respectively. The slides were stored in a dark humid chamber for 30 mins at room temperature. Afterwards, slides were washed again twice for 5 mins and then covered from light with glass coverslips (24 x 50 mm, borosilicate glass, VWR International, Radnor, USA) and VectaMount (Vector Laboratories, USA).

Imaging of H&E and Immunohistology stained slides was performed in the Nikon Imaging Centre Heidelberg, using a Nikon Eclipse Ni-E, equipped with a Nikon DS-Ri2 camera for HE- and a DS-Qi2 camera for Immunofluorescent staining.

## Author contributions

**Concept and study design**: PF, FP, LM, MAC, DM, **Data acquisition**: LM, FP, MAC, DM, MRAPS, **Data analysis**: LM, FP, MAC, **Electron microscopy**: MT, RC, IW, RRS, ***In vivo* study**: LG, NJ, MH, **Writing - original draft preparation**: LM, FP, DM, MAC, MH, **Writing - review and editing**: PF, RRS, MH, **Supervision**: PF, MAC, DM

## Supporting Information

Supporting Information is available from the Wiley Online Library or from the author.

## Acknowledgments

The authors thank Miguel A. Ramos Docampo and Brigitte M. Städler from the Interdisciplinary Nanoscience Center (INANO) in Aarhus Denmark for advice on polymer synthesis. We also thank Eldin Schmitt, Aaron Eidenmüller and Felix Hecht for their support with the dissection of the pig eyes. We would like to thank the Soft (bio)materials characterization Core Facility (Biomechanics) at IMSEAM Heidelberg University, funded by the Federal Ministry of Education and Research (BMBF) and the Ministry of Science Baden-Württemberg within the framework of the Excellence Strategy of the Federal and State Governments of Germany for access to the Cryotome Cryostar NX50. We further thank Bryan C. Ackermann and Gerd U. Auffarth for their support during animal handling. LM, FP, and PF thank Heidi Mühl, José Hurst and Sven Schnichels from the Institute for ophthalmic research, University of Tübingen, Germany, for helpful suggestions. MH is supported by the Add-On Fellowship of the Joachim Herz Foundation. IW, RC, RRS thank the Cluster of Excellence ’3DMM2O’, EXC. LM, FP and PF gratefully acknowledge funding from the *GO-Bio initial* initiative REVeyeVE-2 (grant number 16LW0291) of the Bundesministerium für Bildung und Forschung (BMBF), Germany.

