## Supplementary for "Enzyme-functionalized microparticles to open the vitreoretinal interface"

### Supplementary Information - Eye paper

*Author One  Author Two  Author Three**

A. N. Author, A. N. O. Author
Address
Email Address:
A. N. O. Author
Address

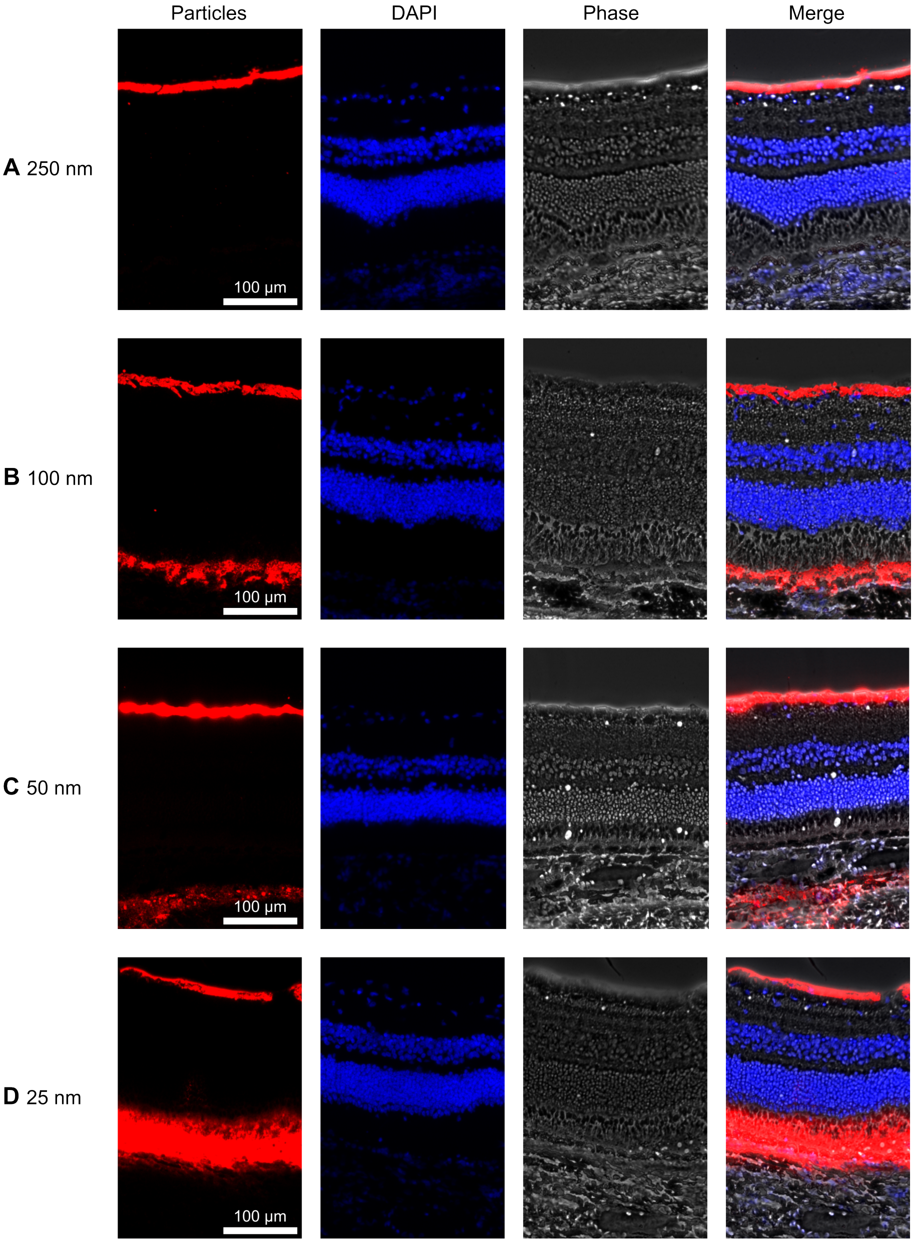


Figure 1: Representative fluorescence images showing retention of fluorescent carboxylated particles at the ILM. Particle sizes (A) 250 nm; (B) 100 nm; (C) 50 nm; and (D) 25 nm. Columns display (from left to right): red fluorescence (nanoparticles), DAPI staining, phase images, and merged images.

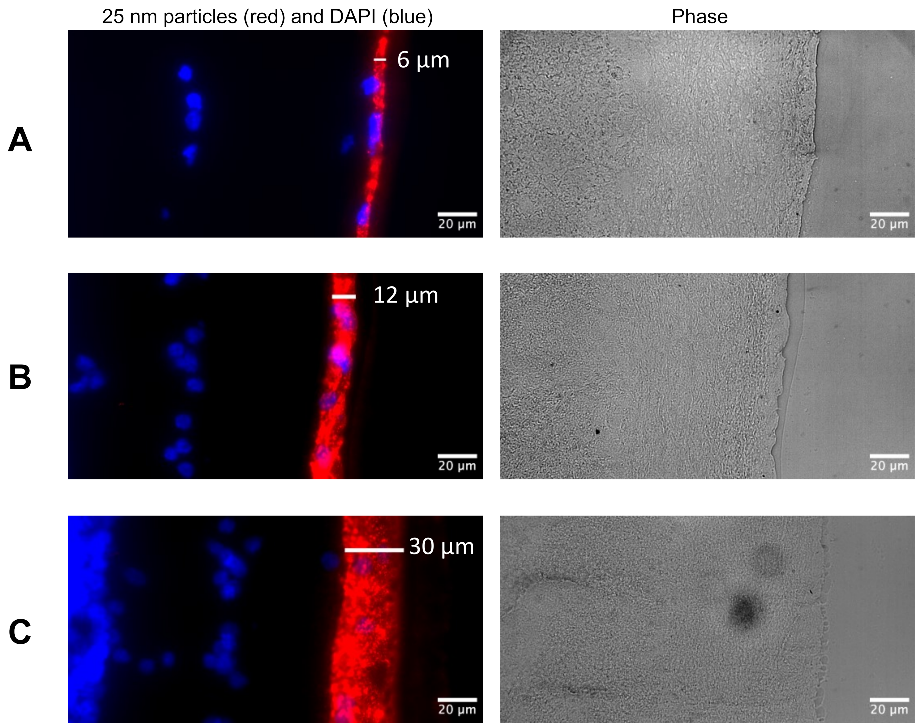


Figure 2: Representative fluorescence images showing the same sample sectioned at varying thicknesses: (A) 6 µm; (B) 12 µm; and (C) 30 µm. Columns display (from left to right): red fluorescence (nanoparticles) together with DAPI staining, and phase images.

#### References
